# supFunSim: spatial filtering toolbox for EEG

**DOI:** 10.1101/618694

**Authors:** Krzysztof Rykaczewski, Jan Nikadon, Włodzisław Duch, Tomasz Piotrowski

## Abstract

Recognition and interpretation of brain activity patterns from EEG or MEG signals is one of the most important tasks in cognitive neuroscience, requiring sophisticated methods of signal processing. The supFunSim library is a new Matlab toolbox which generates accurate EEG forward models and implements a collection of spatial filters for EEG source reconstruction, including linearly constrained minimum-variance (LCMV), eigenspace LCMV, nulling (NL), and minimum-variance pseudo-unbiased reduced-rank (MV-PURE) filters in various versions. It also enables source-level directed connectivity analysis using partial directed coherence (PDC) and directed transfer function (DTF) measures. The supFunSim library is based on the well-known Field-Trip toolbox for EEG and MEG analysis and is written using object-oriented programming paradigm. The resulting modularity of the toolbox enables its simple extensibility. This paper gives a complete overview of the toolbox from both developer and end-user perspectives, including description of the installation process and some use cases.

## 1 Introduction

Network neuroscience is at present the most promising approach to understand the structure and functions of complex brain networks. Neuroimaging and signal processing methods are rapidly evolving, with the ultimate goal of reaching high time and space resolution, allowing for models of functional connectivity, activation of large-scale networks and their rapid dynamic transitions in multiple time scales. Electroencephalography (EEG) has excellent temporal resolution, is noninvasive and relatively easy to use. Unfortunately, signals observed at the scalp level are difficult to interpret, due to the propagation of signals from their cortical and subcortical sources through multiple layers of the brain with several different volume conduction properties, including the scalp, skull, cerebrospinal fluid (CSF), and brain tissues. Sensors receive corrupted mixed signals from various active brain structures. Therefore, direct scalp-level EEG analysis cannot reflect the underlying neurodynamics.

Reconstruction of brain’s electrical activity from electroencephalographic (EEG) or magnetoencephalographic (MEG) recording is frequently based on spatial filters.^1^ Such task requires solving forward and inverse problems in EEG or MEG and is of great interest to EEG/MEG community. Several libraries implementing various forward and inverse solutions are available, see for example [27,37,13,39,12] and references therein. supFunSim extends their functionality by providing object-oriented implementation of both EEG forward and inverse problems. In the former case, it uses realistic and extendable model of activity of sources and enables usage of accurate EEG forward models. In the latter case, it implements a large collection of spatial filters for EEG source reconstruction, including the linearly constrained minimum-variance (LCMV) [10,38,36], eigenspace LCMV [36], nulling (NL) [16], and minimum-variance pseudo-unbiased reduced-rank (MV-PURE) filters [33,30,32]. It also enables source-level directed connectivity analysis using partial directed coherence (PDC) [2] and directed transfer function (DTF) [18] measures that can be applied to the time series representing reconstructed activity of sources of interest.

The supFunSim toolbox is based on FieldTrip [27], an excellent Matlab toolbox for EEG and MEG signal analysis. It can be used as an extension (plugin) to FieldTrip. Thanks to its object-oriented design, functionality of its modules can be easily extended, e.g., by providing data or functions from other toolboxes.

The source code of the current version of the toolbox is publicly available at https://github.com/nikadon/supFunSim.git as an `Org-mode` file, Jupyter notebook, and also as a plain Matlab source code. Functions are not precompiled, as script libraries have the advantage of being easily maintainable and extensible.

The paper is organized as follows. First, we briefly introduce the forward and inverse problems in EEG (similar considerations also apply to MEG). Then, we discuss the benefits of the object-oriented approach for our toolbox and its extensibility. We close with the use cases which appear frequently in practical applications of this toolbox. In appendices mathematical details of the implemented spatial filters, and the full list of toolbox parameters, are provided.

## 2 EEG Measurement Model

Electromagnetic signals that originate within *enchanted loom*^2^ of the human brain are propagated through vaious head compartments [24]. This *dissolving pattern* of brain electrical activity can be detected on the surface of scalp using electroencephalography (EEG).

At a given time instant the EEG data acquisition can be well approximated by a linear equation of the generic form

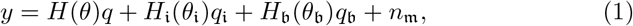

where, for *m* EEG sensors located at the scalp and *l* sources of interest modelled as equivalent current dipoles (ECD) located inside brain’s volume,

– by *y* ∈ ℝ^*m*^ we denote the signal observed in the sensor space at a given time instant,
– *θ* = (*θ*_1_,…,*θ_l_*) ∈ *Θ* ⊂ ℝ^*l*×3^ represents locations of the sources of interest, i.e., for the *i*-th source the vector representing its location is *θ_i_* ∈ ℝ^3^. Here, *Θ* denotes the set of all subsets of locations of source signals,
– *H*(*θ*) ∈ ℝ^*m×l*^ is the sensor array response (*lead-field*) matrix of the sources of interest,
– *q* ∈ ℝ^*l*^ is a vector of electric activity of the *l* sources of interest representing magnitudes of ECDs,
– similarly, for the *k* interfering noise sources, 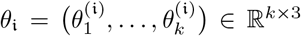 are the interference source locations, 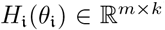 is the corresponding lead-field matrix, 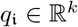 is the corresponding interference activity,
– for the *p* sources of background activity of the brain 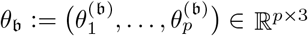 are the background source locations (i.e. sources which are not sources of interest and are uncorrelated with them), 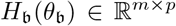 is the corresponding lead-field matrix, 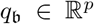 is the corresponding background activity,
– 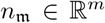 is an additive white Gaussian noise (AWGN) interpreted as a measurement noise present in the sensor space.

Equation (1) represents a single sample of EEG data *y* from subjects’ scalp at a given time (*forward solution*). It enables, through customizable parameters, accurate modelling of real-world EEG experiments. We shall emphasize at this point that not all components of the model in (1) have to be considered, i.e., one may select only those signal components that fit the aims of user’s simulation settings.

The lead-field matrices establishing signal propagation model are estimated on the basis of geometry and electrical conductivity of head compartments together with position of sensors on the scalp. We consider these properties to be fixed in time during a single EEG data acquisition session. Therefore, lead-field-matrices are assumed to be time-invariant in such circumstances (its values do not change during acquisition time in a single session). We also note that the above EEG forward model (1) assumes that the orientations of the ECD moments are fixed during measurement period, and only their magnitudes *q*, 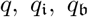 vary in time. We also assume that orientations of the ECD moments are normal and directed outside with respect to the cortical surface mesh. This is in accordance with the widely recognized physiological model of EEG signal origin that considers pyramidal cortical neurons to be the main contributor to the brain’s bioelectrical activity that can be measured on the human scalp [3].

We assume that *q* and 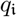 may be correlated (i.e., that sources of interest can interfere with each other), but are uncorrelated with the background sources 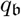 and the noise 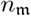. We further assume that *q*, 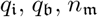 are zero-mean weakly stationary stochastic processes with the exception that *q* may contain in addition a deterministic component simulating evoked (phase-locked) activity in event-related EEG experiments. In our toolbox the presence of this component is controlled by the `SETUP.ERPS` variable.

## 3 EEG Source Reconstruction

Having solved the EEG forward problem which introduced, in particular, the lead-field matrices embodying the propagation model of brain’s electromagnetic activity, we are in a position to solve the inverse problem. Here it amounts to reconstruction of time courses of activity of sources at predefined locations. That means that we assume that the locations of the sources of interest *θ* are known. This can be achieved by defining regions of interest using source localization methods, e.g., minimum-norm [29] or spatial filtering-based methods [23,31], or referring to neuroscience studies that have identified regions of interest (see for example Hui et al. [16]). Then, the goal is to reconstruct the activity *q* of sources of interest based on the observed signal *y* as

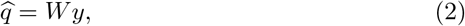

where *W* ∈ ℝ^*l×m*^ is a matrix representing spatial filter’s coefficients. The definitions of the filters currently implemented in the toolbox are given in Appendix *Implemented spatial filters*.

**Fig. 1.**
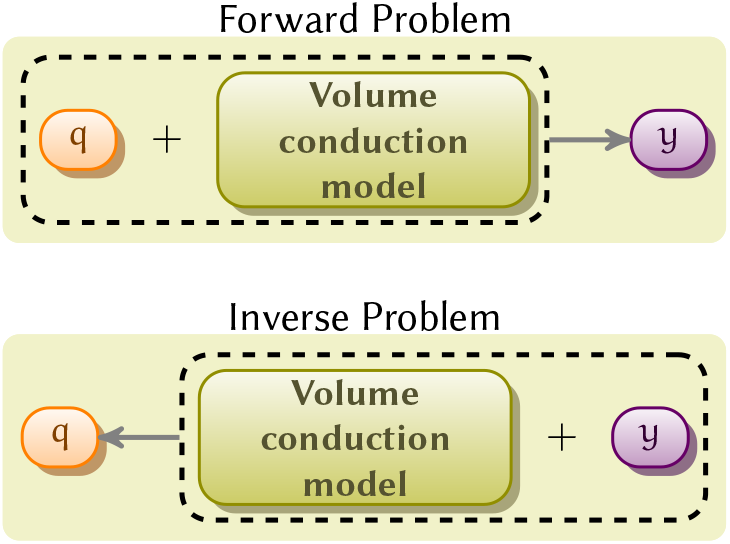
The relationship between forward and inverse problems.

## 4 Toolbox Signal Processing Outline

### 4.1 Overview

In order to obtain EEG signal *y* we need first to generate source activity signals and propagate them to the position of electrodes, according to the forward model (FM) given in Equation (1). The source signals are generated using the method described in [9], which uses stable multivariate autoregression (MVAR) model with predefined coefficient matrices. This results in a wide-sense stationary signals generated with predefined pairwise linear dependencies (correlations). Such approach has been studied and used in literature, see, e.g., [14,2,25], and is especially useful in investigating functional dependencies between activity of sources using directed connectivity measures such as partial directed coherence (PDC) [2] or directed transfer function (DTF) [21].

Gaussian pseudo-random vectors simulating autoregressive processes are generated using the `arsim` function available from [35]. In this way, we obtain multivariate time series representing activity of sources of interest *q* (denoted in the code as `sim_sig_SrcActiv.sigSRc`), sources of interference due to noise 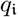 (`sim_sig_IntNoise.sigSRc`), and sources of background activity 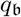 (`sim_sig_BcgNoise.sigSRC`).

Moreover, as our framework allows to add event-related potentials (ERP, flag variable `SETUP.ERPs` in the code) to the source signal, each source activity is divided into *pre* and *pst* parts in relation to the onset of event (*q* into *q_pre_* and *q_pst_*, etc.) in order to enable simulation of ERP experiments. In particular, the ERP signal may be added to *q_pst_*, but not to *q_pre_*.

Furthermore, the *pre* and *pst* subsignals are used to implement spatial filters. In particular, noise correlation matrix *N* may be estimated from *y_pre_* signal and signal correlation matrix *R* may be estimated from *y_pst_* signal.

The *y_pst_* signal is also used for evaluation of the fidelity of reconstruction, as a filter *W_f_* produces an estimate of the activity signal source 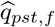 based on *y_pst_*. Then, the MVAR model, which is represented by composite model matrix A00, is fitted to the reconstructed source activity using arfit function, yielding reconstructed composite MVAR model matrix A00_*f*_. This matrix can then be used to investigate directed connectivity using partial directed coherence (PDC) [2] and directed transfer function (DTF) [18].

The overview of signals processed by the toolbox and dependencies between them is depicted in Figure 2.

**Fig. 2.**
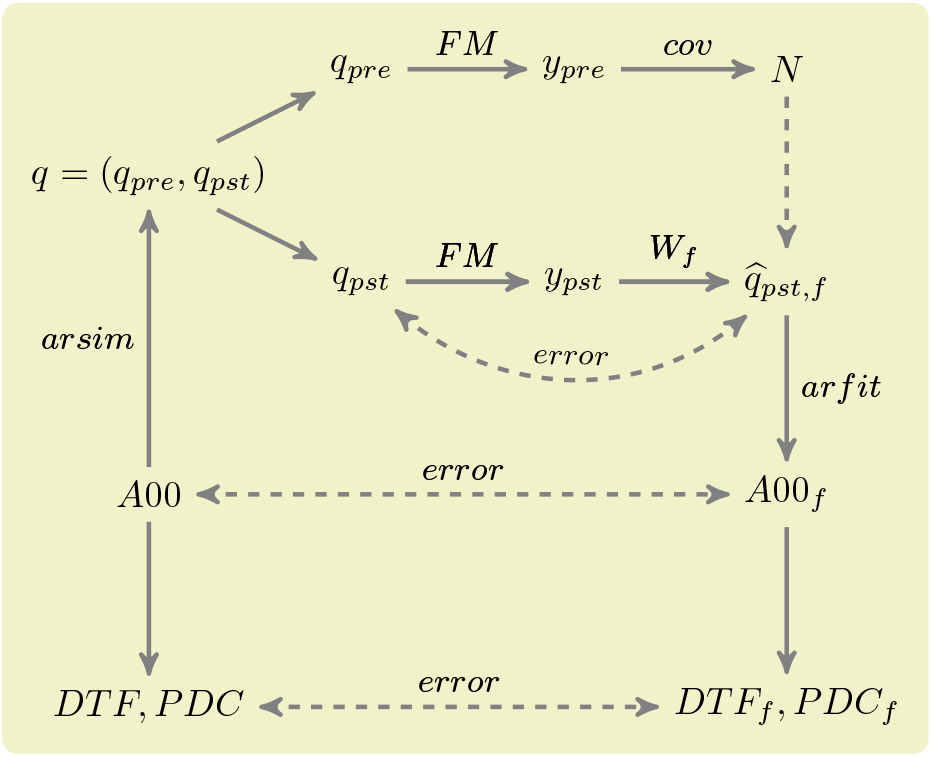
Overview of processing of signals by out toolbox. `arsim` and `arfit` functions are taken from [35], *A*00_*f*_ is the reconstructed composite MVAR model matrix, and *W_f_* is the matrix with spatial filter’s coefficients.

### 4.2 Brain Signals in the Source Space

Items 1 and 2 below describe generation of source signals *q*, 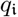 and 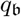. Item 3 concerns definition of volume conduction model. The computation of lead-field matrices *H*, 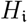 and 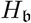 is discussed in the subsequent Section *Brain Signals in Sensor Space*.

#### 1. Positioning sources

File `supFunSim/mat/sel_atl.mat` contains 15000 vertex FreeSurfer [5] brain tessellation together with atlases [5,8] that provide parcellation of the mesh elements into cortical patches (regions of interest, ROIs). This file is provided with the BrainStorm toolbox [37]. First, we randomly select an arbitrary number of ROIs by choosing items from Destrieux and Desikan-Killiany atlases [8,7]. In each ROI, each vertex is a candidate node for location of the dipole source. Then, an arbitrary number of locations can be drawn within each ROI, separately for *q*, 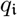 and 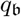.

Most of the simulation parameters are controlled using `SETUP` structure. The geometrical arrangement and number of cortical sources in each ROIs is controlled using `SRCS` field (`SETUP.SRCS`), which is a three-column `<int>` array, where:

– rows represent consequent ROIs (thus, the number of rows determines the number of ROIs used in the simulations),
– the first column represents sources of interest, the second column represents sources of interference, and the third column represents sources of background activity,
– integer values of this array represent number of sources in the given ROI for the given signal type.

For the end-user this provides mechanism not only to control the total number of sources of a particular type, but also to choose their spatial distribution. Additionally, we provide a mesh representing both *thalami* (jointly) as a structure containing potential candidates for the non-cortical (deep) sources of signal/noise. Variable `SETUP` contains also the field (`SETUP.DEEP`) which defines the number of signals arising in the brain center (around thalami) belonging to a particular signal type (of interest, interference, or background activity). Furthermore, in order to account for the mislocalization of sources, together with the original lead-fields we also generate *perturbed* lead-fields for the source activity reconstruction. These are generated using locations that are shifted with respect to the original locations in the direction that is rotated in relation to the original (normal to cortex surface) dipole orientation. Default shift is random and less then 5 mm in each direction (*x, y, z*). Default rotation is random, bounded by 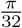 (azimuth and elevation) angle.

#### 2. Sources Timecourse

Following [25,35], we use stable autoregressive (MVAR) model to generate time-series. It is assumed that such model generates a realistic source activity [20]. We create separate models for time-series *q* and 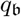. The 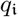 is obtained as a negative of *q* with Gaussian uncorrelated noise added with the same power as the *q*, i.e. 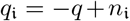. In this way, we obtain correlated time-series *q* and 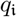.

The *l*-variate autoregressive model of order *p* for a stationary time-series of state vectors *q^n^* ∈ ℝ^*l*^ is defined at time instant *n* as

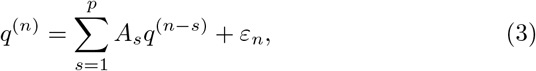

where *q*^(*n*)^ is the state vector at time *n, p* is the order of the model (*p* = 6 by default), matrices *A*_1_,…,*A_p_* ∈ ℝ^*l×l*^ are the coefficient matrices of the AR model, and *ε_n_* is the *l*-dimensional additive white Gaussian noise [14]. For the signal of interest *q*, we also give the possibility to include deterministic component simulating evoked (phase-locked) activity in event-related EEG experiments. The presence of this component is controlled by the `SETUP.ERPs` variable. Then, *q* = *q*^(*n*)^ + *q*^(*d*)^, where *q*^(*d*)^ is the deterministic ERP component. In the toolbox, *q*^(*d*)^ is generated using Matlab `gauswavf` function generating first derivative of Gaussian wavelet function, as recommended by [34].

Similarly, 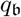 is simulated using independent, random and stable MVAR model (of order *r* = 6 by default):

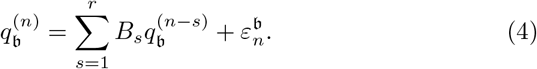

MVAR model is considered to be *stable* if the absolute values of all eigenvalues of all matrices *A_s_* (respectively, *B_s_*) are less than one. We used procedure adapted from [11] (namely, we adapted the `stablemvar` function) to generate stable MVAR model that was used for times-series generation. During generation of MVAR models for *q* and 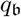, the coefficient matrix *A_s_* (respectively, *B_s_*) is multiplied by a masking matrix that has 80% (by default) of its off-diagonal elements equal to zero. All the remaining diagonal and off-diagonal masking coefficients are equal to one. In code, the composite MVAR model matrix is represented by the variable A00, see Fig. 3. Such procedure allows, in particular, to obtain specific profile of directed dependencies between activity of sources of interest. This approach is taken from [2]. Moreover, it gives us the possibility to implement the PDC and DTF directional casual dependency measures. Namely, partial directed coherence and directed transfer function are matrices defined using Fourier transform of MVAR model (3), i.e.

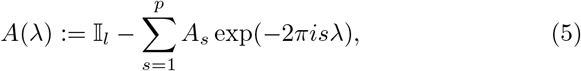

where 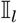 is the identity matrix, λ is normalized frequency, |λ| ≤ 0.5. Then, partial directed coherence between *i*-th and *j*-th signals is given by [2]

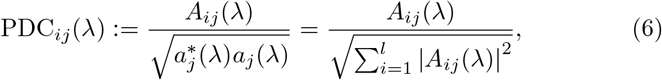

where *A_ij_*(λ) is *ij* element of matrix *A*(λ), *a_j_*(λ) is *j*th column of *A*(λ) and * means Hermitian transpose. It takes values in the interval [0,1] and measures the relative strength of the interaction of a given source signal *j* to source signal *i* normalized by strength of all of *j*’s connections to other signals [4].

**Fig. 3.**
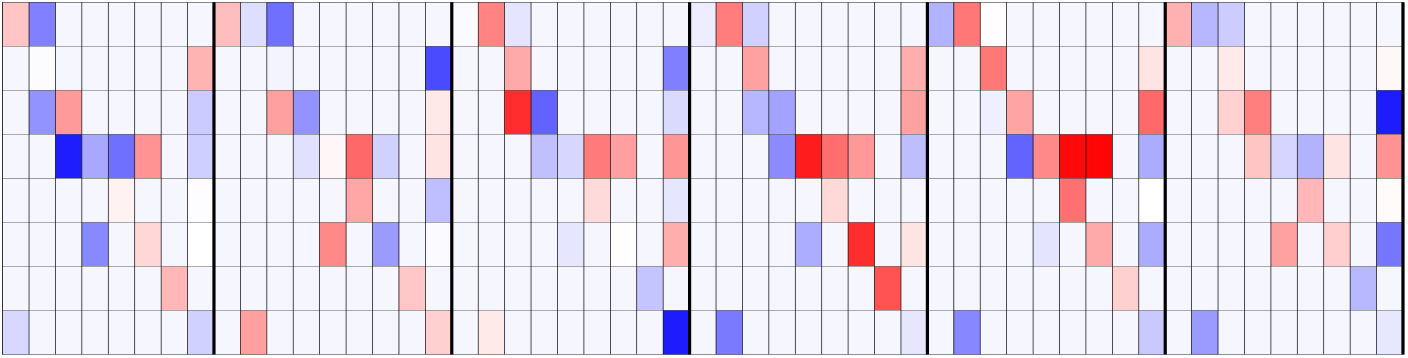
*A*00: the original coefficient matrix used for time-series generation, with sample values *l* = 9 and *p* = 6., after application of a random mask.

Directed Transfer Function (DTF) is defined as

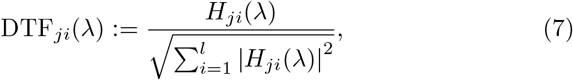

where *H*(λ):= (*I* − *A*(λ))^−1^ is the **transfer matrix**, *i, j* = 1,…,*l*. It can be interpreted as ratio between inflow from channel *i* to channel *j* normalized by the sum of inflows to channel *j*.

#### 3. Volume conduction model

We used FieldTrip toolbox [28] to generate volume conduction model (VCM) and lead-field matrices. VCM was prepared using FieldTrip `ft_prepare_headmodel` function implementing `DIPOLI` method [26], which takes as arguments three triangulated surface meshes representing the outer surfaces of brain, skull and `scalp supFunSim/mat/msh.mat` [37]. VCM is available in our toolbox in the precomputed form (`supFunSim/mat/sel_vol.mat`), although, if required, FieldTrip toolbox allows for easy computation of custom VCMs on the basis of triangulated meshes which can be obtained from structural (T1) MRI scans. Thanks to the modular design of our toolbox other VCMs may also be easily imported.

The default head geometry is based on the Colin27^3^ [37,15,1]. However, it can be easily substituted at user’s discretion by replacing triangulation meshes stored in `SETUP.sel_msh` (a list of structures containing: scalp outer mesh, skull outer mesh and skull inner mesh, where the last triangulation represents “rough” brain outer mesh). Common choices include realistic head models generated on the basis of structural MRI scans or spherical models.

### 4.3 Brain Signals in the Sensor Space

In the sensor space we need to provide positions for the electrodes (*a.k.a*. sensor montage). By default, in our simulations we use *HydroCel Geodesic Sensor Net* utilizing 128 channels as EEG cap layout. Other caps can easily be used by substituting content of the `supFunSim/mat/sel_ele.mat` with electrode position coordinates obtained either from specific EEG cap producer, or from the standard montages that are available with EEG analysis software such as EEGLAB [6] or FieldTrip [28]. Additionally, for real data acquisition setup, the electrode positions can be captured using specialized tracking system for every EEG session.

The volume conduction model, together with source locations and their orientations, are obtained as described in the three points of the previous section. Together with electrode positions, they are the input arguments for the `ft_prepare_leadfield` function, which is run three times during simulations, outputting *H*, 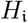 and 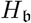.

## 5 Implementation Details

### 5.1 Object-oriented approach

The object-oriented approach provides the toolbox with several desirable properties of the code and avoids drawbacks of standard procedural approach commonly employed in Matlab scripts. For example, Matlab by default stores all variables in one common workspace. This causes bugs in the code that may be hard to detect. On the other hand, the object-oriented approach circumvents this difficulty by its inherent encapsulation property, enclosing variables within a class, and sealing it securely from the outside environment. We also note that the construction of the Matlab language requires explicit assignment of an instance of the class each time a method acts on it. This approach necessitates language constructs such as `obj = obj.method`, where `obj` is a given instance of a class.

Data structures created during simulation can be accessed interactively in the Jupyter notebook or in the Matlab script. In particular, property `MODEL` from `EEGReconstruction` class contains information about all variables used within the simulation pipeline.

### 5.2 Benefits of literate programming

The code of the toolbox was written in Jupyter, which is an open source application that allows users to create interactive and shareable notebooks. Jupyter allows for easy export of whole documents to HTML, LATEX, PDF and other formats, and is a very convenient tool for academic prototyping, because it permits comments in the code using LATEX mathematical expressions. The source code blocks are interspersed with ordinary natural language blocks that provide explanations and some insights explaining the intrinsic mechanics of the code. Such an approach is called *literate programming* [19]. An example of mathematical comment and corresponding code is given in Figure 4.

**Fig. 4.**
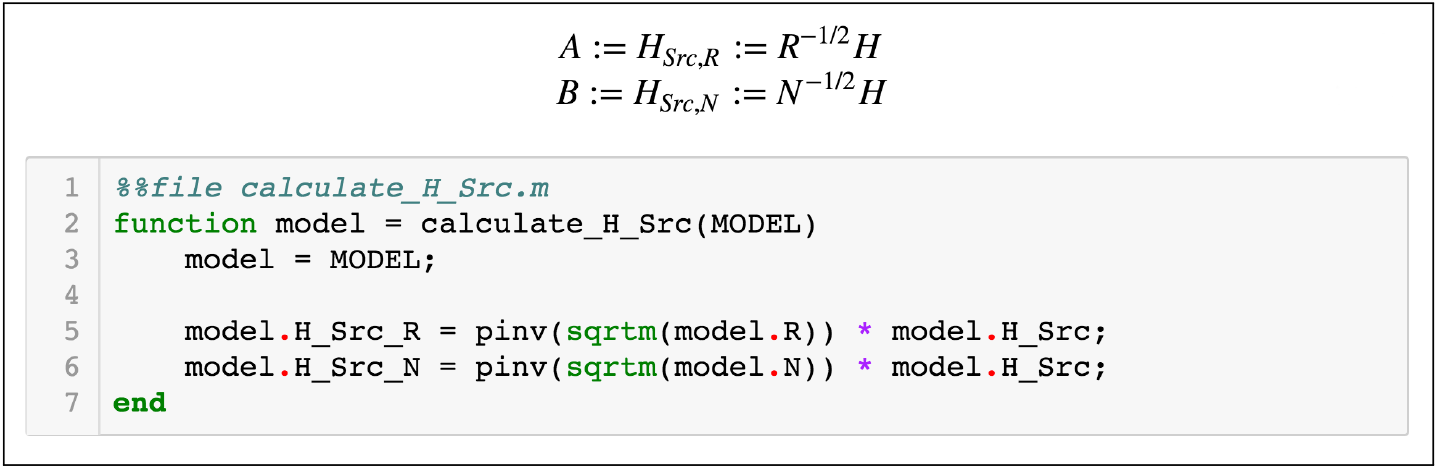
Example of Jupyter notebook.

However, the use of Jupyter is not necessary to run the toolbox. Instead, the code can be executed under powerful and cross-platform Matlab environment. To that end we have prepared a version of the toolbox as the set of Matlab files stored in `supFunSim.zip` archive.

The toolbox does not have a GUI (Graphical User Interface). Instead, user interacts with it using provided functions. Therefore, as a prerequisite to use the supFunSim toolbox knowledge of Matlab language basics is required.

### 5.3 Installation

Installation of the supFunSim is independent of the operating system. For a simple installation similar to FieldTrip’s installation process, the user can download the file `supFunSim.zip` from https://github.com/nikadon/supFunSim.git. This archive contains the whole toolbox. After unpacking this archive, the user should execute `addpath(genpath(‘/path/to/toolboxes/supFunSim/’]))`. Function genpath will ensure that all subdirectories will be added to your path. It is most convenient to have the `addpath` function in the `startup.m` script located in the Matlab directory. Then, the user may run the `RunAll.m` script (preferably line by line, in order to follow execution). The user has to make sure that there is a `mat/` directory (or a link to it) containing `mat` files required by the toolbox in the toolbox directory. The `mat` files are available for download at http://fizyka.umk.pl/~tpiotrowski/supFunSim.

More advanced user may manipulate Jupyter notebooks directly and use `make` tool to set up the toolbox from scratch. Namely, in order to open and run notebooks the user should download and install Jupyter notebook with Matlab kernel. The easiest way to do it (under a Unix-like system) is by executing the following instructions in the command line. First, we set up a virtual environment, which will install Python packages locally:

~~~
1 sudo pip install virtualenv # installing virtualenv environment
2 mkdir supFunSimToolbox # making directory for virtual environment
3 unzip supFunSim.zip -d supFunSimToolbox # extracting toolbox
4 virtualenv supFunSimToolbox # creating virtual environment
5 source./bin/activate # activating virtual environment
~~~

Next, we install all necessary packages and install Matlab Engine API for Python

~~~
1 pip install -r requirements.txt # installing all requirements
2 cd /path/to/matlabroot/extern/engines/python
3 python setup.py install
~~~

We also provide a `make` tool for a simple administration of notebooks’ code. For example, the user may execute `make everything` in terminal in order to generate all source code files. See `README.md` file in the repository for details.

At this stage, one can run the simulations and “play with” the code by going to `supFunSimToolbox` directory and running

~~~
1 jupyter-notebook
~~~

Finally, the description of the installation under Windows can be found in the `README.md` file.

### 5.4 Prerequisites/dependencies

In addition to Matlab our toolbox requires 3 packages: FieldTrip toolbox (version at least 20150227) [27], MVARICA toolbox (version at least 20080323) [11], ARfit toolbox (version at least 20060713) [25,35]. Location of these toolboxes should be added to Matlab `path`.

## 6 Application structure

The simulation framework provided with the current paper consists of a set of modules represented by corresponding classes. The classes are defined in separate (self-contained) notebooks. The classes depend on auxiliary functions generated alongside with them when appropriate `make` target is invoked. In this way, a given class is enclosed and all operations involving it are made within it. The toolbox contains six classes (described in the next section) with a number of auxiliary functions.

### 6.1 Overview of toolbox classes

The main functionality of the toolbox is provided by the following five classes:

– `EEGParameters.ipynb` — class generating parameters for simulations. It can be overwritten in order to obtain desired parameters for a sequence of simulations.
– `EEGSignalGenerator.ipynb` — class used to generate signal for forward modelling of sources. It can be overwritten to generate a signal with given or desired properties.
– `EEGForwardModel.ipynb` — class implementing forward model. It constructs and adds together all signals (source activity, background activity and interference noise). Furthermore, the lead-field matrix is build using FieldTrip library based on the selected head model.
– `EEGReconstruction.ipynb` — class implementing methods used in the reconstruction of the underlying neuronal activity. All spatial filters are implemented in this class.
– `EEGPlotting.ipynb` — class implementing plots detailing execution of experiments. Various visualizations are accessible through the methods included in this class.

We also wrote a class for unit testing of the toolbox functionality:

– `EEGTest.ipynb` — class implementing unit tests and validation of the code against the functional-code toolbox implementation.

Fig. 5 gives an overview of relationships between implemented classes. We should also emphasize that modular design facilitates reuse and extensibility of the source code and adaptation to other applications. For example, if one wishes to generate source signals in a different way compared with the implemented version, one needs only to overwrite `EEGSignalGenerator` class (or some of its methods) while keeping the rest of the code and its functionality intact.

**Fig. 5.**
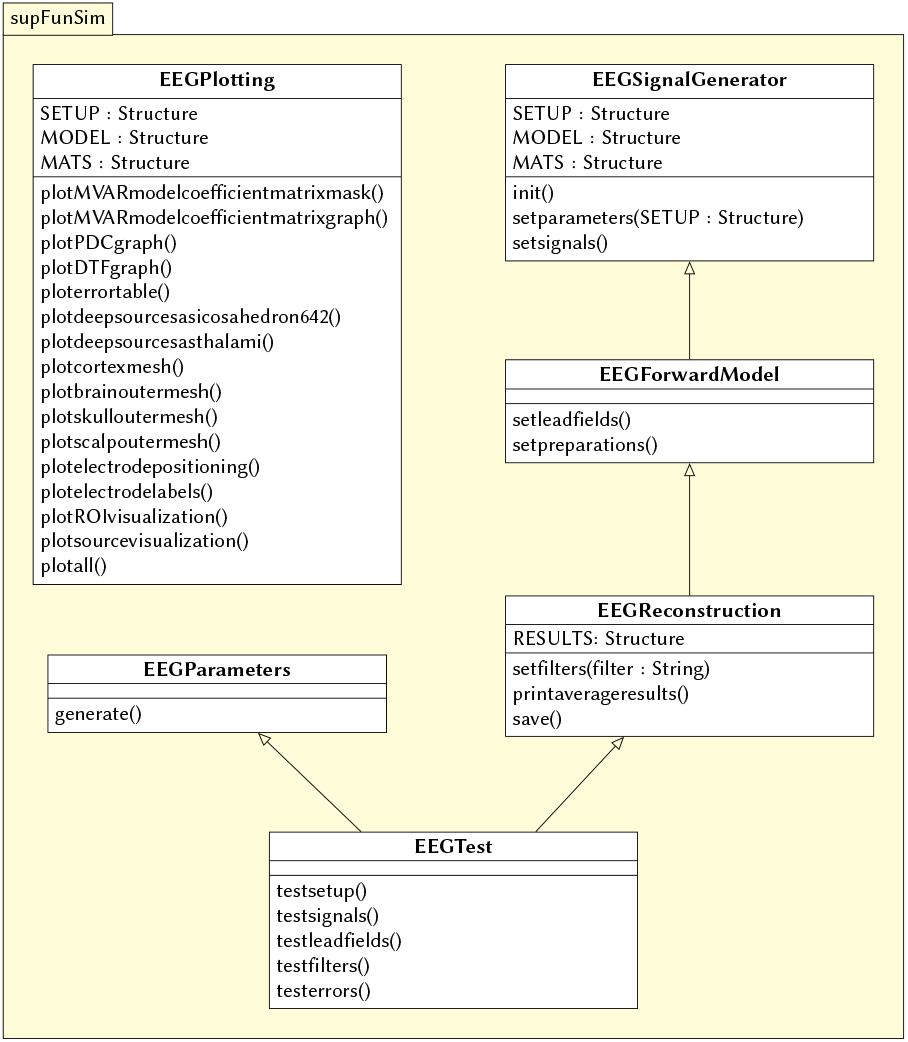
Dependencies between classes.

### 6.2 Mat files

Directory `mat/` contains third-party data with the geometry of the brain, taken from the Brainstorm toolbox [37] and extracted using FieldTrip procedures:

– `sel_msh.mat` — head compartments geometry (vertices and triangulation forming meshes for brain, skull and scalp); this data can be used as an input for volume conduction model and lead-field generation using FieldTrip (or any other toolbox that can be used to generate forward model);
– `sel_vol.mat` — volume conduction model (head-model). This structure contains head compartments geometry (the same as in `sel_msh.mat`) accompanied by their conductivity values and a matrix containing numerical solution (utilizing boundary or finite element method) to a system of differential equations describing propagation of the electric field. This data is obtained using FieldTrip’s `dipoli` method and is used as an input to the function that calculates the lead-field matrix. The default volume conduction model was prepared in accordance with the instruction provided in the FieldTrip tutorial *Creating a BEM volume conduction model of the head for source-reconstruction of EEG data*.^4^ This structure is used to compute lead-fields;
– `sel_geo_deep_thalami.mat` — mesh containing candidates for location of deep sources (based on *thalami*). The mesh was prepared on the basis of the Colin27 [37,15,1] MRI images;
– `sel_geo_deep_icosahedron642.mat` — mesh containing candidates for location of deep sources (based on *icosahedron642*);
– `sel_atl.mat` — cortex geometry with (anatomical) ROI parcellation (cortex atlas). This detailed triangulation is parceled into cortical patches (a.k.a. regions of interest, ROIs). It contains a 15000 vertices and it is based on the sample data accompanying the BrainStorm toolbox [37]. It was originally prepared using FreeSurfer [5, 8] software;
– `sel_ele.mat` — geometry of electrode positions. By default we use *HydroCel Geodesic Sensor Net* sensor montage utilizing 128 channels available. The electrode positions file is available with the FieldTrip toolbox as `GSN-HydroCel-128.sfp file`;
– `sel_src.mat` — lead-fields of all possible source locations.

#### 6.2.1 Simulation parameters class

This class is responsible for setting up parameters of simulations.

– `EEGParameters:`

– `generate` — this method generates the set of parameters for simulations. The method itself is mainly based on `generatedummysetup` function which itself uses `setinitialvalues` and `setsnrvalues` functions containing default configuration for the reconstruction. Users willing to change basic configuration should edit the `configurationparameters.m` file. The assignment of parameters’ values made in this file overwrites default parameter settings. For the complete list of all simulation parameters consult Table 1. For unit testing, configuration from the `testparameters` should be used. The configuration in this file agrees with the configuration used in the `supFunSim.org` file. To perform unit testing, simply uncomment appropriate line in the `generatedummysetup.m` file.

**Table 1.** `SETUP` configuration.

| Parameter | Description |
| --- | --- |
| `rROI` | random (if 1) or predefined (if 0) ROIs |
| `rPNT` | random (if 1) or predefined (if 0) candidate points for source locations |
| `SRCS` | represent `SrcActiv`, `IntNoise` and `BcgNoise`, respectively |
| `DEEP` | deep sources |
| `ERPs` | add ERPs (timelocked activity) |
| `n00` | number of time samples per trial |
| `K00` | number of independent realizations of signal and noise based on generated MVAR model |
| `P00` | order of the MVAR model used to generate time-courses for signal of interest |
| `FRAC` | proportion of ones to zeros in off-diagonal elements of the MVAR coefficients masking array |
| `STAB` | VAR stability limit for MVAR eigenvalues (less than 1.0 results in more stable model producing more stationary signals [25]) |
| `RNG` | range for pseudo-random sampling of eigenvalues for MVAR coefficients range |
| `ITER` | iterations limit for MVAR pseudo-random sampling and stability verification |
| `PDC_RES` | frequency resolution vector for normalized PDC and DTF estimation |
| `TELL` | provide additional comments during code execution (“tell me more”) |
| `PLOT` | plot figures during the intermediate stages |
| `SCRN` | get screens positions |
| `DISP` | force figures to be displayed on (3dr) screen |
| `SEED` | seed for random number generation |
| `SEEDS` | hard-coded seeds to ensure repeatability of the simulation |
| `RANK_EIG` | rank of EIG-LCMV filter: set to number of active sources |
| `fltREMOVE` | to keep (if 0) or remove (if 1) selected filters |
| **SHOWori** | to show (if 1) or do not show (if 0) Original and Dummy signals on figures |
| **IntLfgRANK** | rank of patch-constrained reduced-rank lead-field |
| **supSwitch** | **rec**: run reconstruction of sources activity, **loc**: find active sources |
| **thalamus** | type of head model |
| **DEBUG** | if we want to debug |
| **PATH** | path to directory with the code |
| **SRATE** | sampling rate |
| **CUBE** | perturbation of the lead-fields based on the shift of source location within a cube of given edge length (centered at the original lead-fields locations) |
| **CONE** | perturbation of the lead-fields based on the rotation of source orientation (azimuth TH, elevation PHI) |
| **H_Src_pert** | use original (if 0) or perturbed (if 1) lead-field for signal reconstruction |
| **H_Int_pert** | use original (if 0) or perturbed (if 1) lead-field for nulling constrains |
| **SINR** | signal to interference noise power ratio expressed in dB (both measured on electrode level) |
| **SBNR** | signal to biological noise power ratio expressed in dB (both measured on electrode level) |
| **SMNR** | signal to measurment noise power ratio expressed in dB (both measured on electrode level) |
| **WhtNoiseAddFlg** | white noise admixture in biological noise interference noise (FLAG) |
| **WhtNoiseAddSNR** | SNR of **BcgNoise** and **WhiNo** (dB) |
| **SigPre** | final signal components for pre-interval (use zero or one for signal) |
| **IntPre** | final signal components for pre-interval (use zero or one for interference noise) |
| **BcgPre** | final signal components for pre-interval (use zero or one for background activity) |
| **MesPre** | final signal components for pre-interval (use zero or one for measurement noise) |
| **SigPst** | final signal components for post-interval (use zero or one for signal) |
| **IntPst** | final signal components for post-interval (use zero or one for interference noise) |
| **BcgPst** | final signal components for post-interval (use zero or one for background activity) |
| **MesPst** | final signal components for post-interval (use zero or one for measurement noise) |
| **DATE** | date |
| **NAME** | temporary file name |
| **SINR_RNG** | range of SNR for interference signals |
| **SBNR_RNG** | range of SNR for background signals |
| **SMNR_RNG** | range of SNR for measurment noise |

#### 6.2.2 Signal generation class

This class is responsible for EEG signal generation.

– `EEGSignalGenerator:`

– `init` — this method initializes all toolboxes required by supFunSim. It sets path to toolboxes, creates their default settings, sets head model, geometry of patches etc.;
– `setparameters` — this method sets configuration for simulation using parameters from `EEGParameters` class;
– `setsignals` — this method generates source-level signals: `sim_sig_SrcActiv.sigSRC, sim_sig_IntNoise.sigSRC, sim_sig_BcgNoise.sigSRC`, as well as sensor level `sim_sig_MesNoise.sigSNS` measurement noise:

- `makeSimSig` — MVAR-based signal generation; the signal is then divided into *pre* and *pst* parts,
- `generatetimeseriessourceactivity` — generates signals of sources of interest `sim_sig_Src_Activ.sigSRC` using `makeSimSig` and if required, adds ERP deterministic signal to the *pst* part of the signal of interest,
- `generatetimeseriesinterferencenoise` — generates interference noise signal sim_sig_IntNoise.sigSRC as the negative of `sim_sig_SrcActiv.sigSRC` signal of interest with added white Gaussian noise of prescribed power relative to the power of `sim_sig_SrcActiv.sigSRC`,
- `generatetimeseriesbackgroundnoise` — generates background activity signal `sim_sig_BcgNoise.sigSRC` using `makeSimSig`,
- `generatetimeseriesmeasurementnoise` — generates measurement (sensor-level) noise signal `sim_sig_MesNoise.sigSNS` as an additive white Gaussian noise. Class EEGForwardModel generates solution to the forward problem: leadfield matrices and the resulting electrode-level signal.

#### 6.2.3 Forward model class

Class `EEGForwardModel` generates solution to the forward problem: leadfield matrices and the resulting electrode-level signal.

– `EEGForwardModel:`

– `setleadfields` — this method generates leadfield matrices:

- `geometryrandomsampling` — random or user-defined selection of cortex ROIs for sources (of interest, interfering activity, background activity),
- `geometryindices` — identification of cortex ROIs’ indices within cortex atlas,
- `geometrycoordinates` — coordinates of vertices of sources within selected ROIs,
- `geometrydeepsources` — coordinates of vertices of sources within thalamus,
- `geometryperturbation` — generation of perturbed source locations and orientations,
- `geometryleadfieldscomputation` — computation of original lead-fields of sources of interest `sim_lfg_SrcActiv_orig.LFG`, sources of interfering activity `sim_lfg_IntNoise_orig.LFG`, sources of background activity `sim_lfg_BcgNoise_orig.LFG`, as well as their respecitve perturbed versions `sim_lfg_SrcActiv_pert.LFG, sim_lfg_IntNoise_pert.LFG`, and `sim_lfg_BcgNoise_pert.LFG`, respectively,
- `forwardmodeling` — multiplication of source signals by their corresponding lead-field matrices yielding sensor-level signals of sources of interest `sim_sig_SrcActiv.sigSNS`, sources of interfering activity `sim_sig_IntNoise.sigSNS`, and sources of background activity `sim_sig_BcgNoise.sigSNS`;
– `setpreparations` — generates output EEG signal and prepares for signal reconstruction using spatial filters:

- `preparationsnrsadjustment` — rescales sensor-level signals to the appropriate SNRs,
- `prepareleadfieldconfiguration` — determines whether original or perturbed lead-field matrices of sources of interest and of interfering sources will be made available to spatial filters according to the user’s setting of `SETUP.H_Src_pert` and `SETUP.H_Int_pert` flags; moreover, reduces the rank of lead-field matrix of interfering sources according to `SETUP.IntLfgRANK` variable value,
- `preparemeasuredsignals` — sums sensor-level signals and adds measurement noise signal (`sim_sig_MesNoise.sigSNS`) to produce output `y_Pre` and `y_Pst` EEG signals.
- `store2eeglab` — a method that allows the user to save data in the EEGLAB format; this toolbox and its plugins allows for better preprocessing and visualization of EEG signal.
- `rawAdjTotSNRdB` — adjusts `sim_sig_IntNoise.sigSNS, sim_sig_BcgNoise.sigSNS` and `sim_sig_MesNoise.sigSNS` signals’ power with respect to the power of `sim_sig_SrcActiv.sigSNS`, to obtained desired signal-to-interference, signal-to-background-activity, and signal-to-noise ratios, respectively. These ratios are defined using `SETUP.SINR, SETUP.SBNR`, and `SETUP.SMNR` variables, respectively, and are expressed in decibels [dB] using the following implementation:

~~~
1 function [y] = rawAdjTotSNRdB(x01, x02, newSNR)
2   y = ((x02 / norm(x02)) * norm(x01)) / (db2pow(0.5 * newSNR)),
3 end
~~~

where `db2pow` is the Matlab function converting decibels to power.

#### 6.2.4 Reconstruction class

Class `EEGReconstruction` computes implemented spatial filters and applies them to the observed `y_Pst` simulated EEG signal. It also computes various fidelity measures of the reconstruced activity of sources of interest.

– `EEGReconstruction:`

– `setfilters` — calculates matrices which are used in the process of reconstruction:

- `spatialfilterconstants` — We compute some of constants used later in defining filters.
- `spatialfiltering` — We compute all intermediate variables needed to calculate the filters.
- `spatialfilteringexecution` — For every filter *W*(*θ*) given as a parameter to this function we are calculating *W*(*θ*)*y* using post-interval signal as *y*. Then we perform arfit for all reconstructed signals and obtain autoregression matrix. This matrix is necessary to calculate PDC and DTF measures.
- `spatialfilteringerrorevaluation` — Function which calculates difference between original and reconstructed signals. It uses various measures: Euclidean metric and correlation coefficients to compare activity signals, MVAR coefficient matrices and PDC and DTF coefficient matrices.
- `vectorizerrorevaluation` — Function that combines results in a single array. Such uniform output of different error measures is later used in plotting.
– `printaverageresults` — print table of comparison of different reconstruction filters.
– `save` — save reconstructed filters.

### 6.3 Plotting class

The toolbox makes it possible to visualize results of experiments. User can plot results of simulation using `EEGPlotting` class which is specially prepared for this.

~~~
1 eegplot = EEGPlotting(reconstruction);
~~~

Interesingly, there are many ways to facilitates visualization of the analysis and results. Jupyter functionality gives us a possibility to plot figures inside notebook (using `%plot` magic option):

~~~
1 %plot inline
2 eegplot.plotskulloutermesh();
~~~

or to open it in Matlab interactive environment:

~~~
1 %plot native
2 eegplot.plotsourcevisualization();
~~~

Toolbox supFunSim contains variety of plotting functions for different visualizations of results. Some of them are presented in Figures 6 7 8 and 9.

**Fig. 6.**
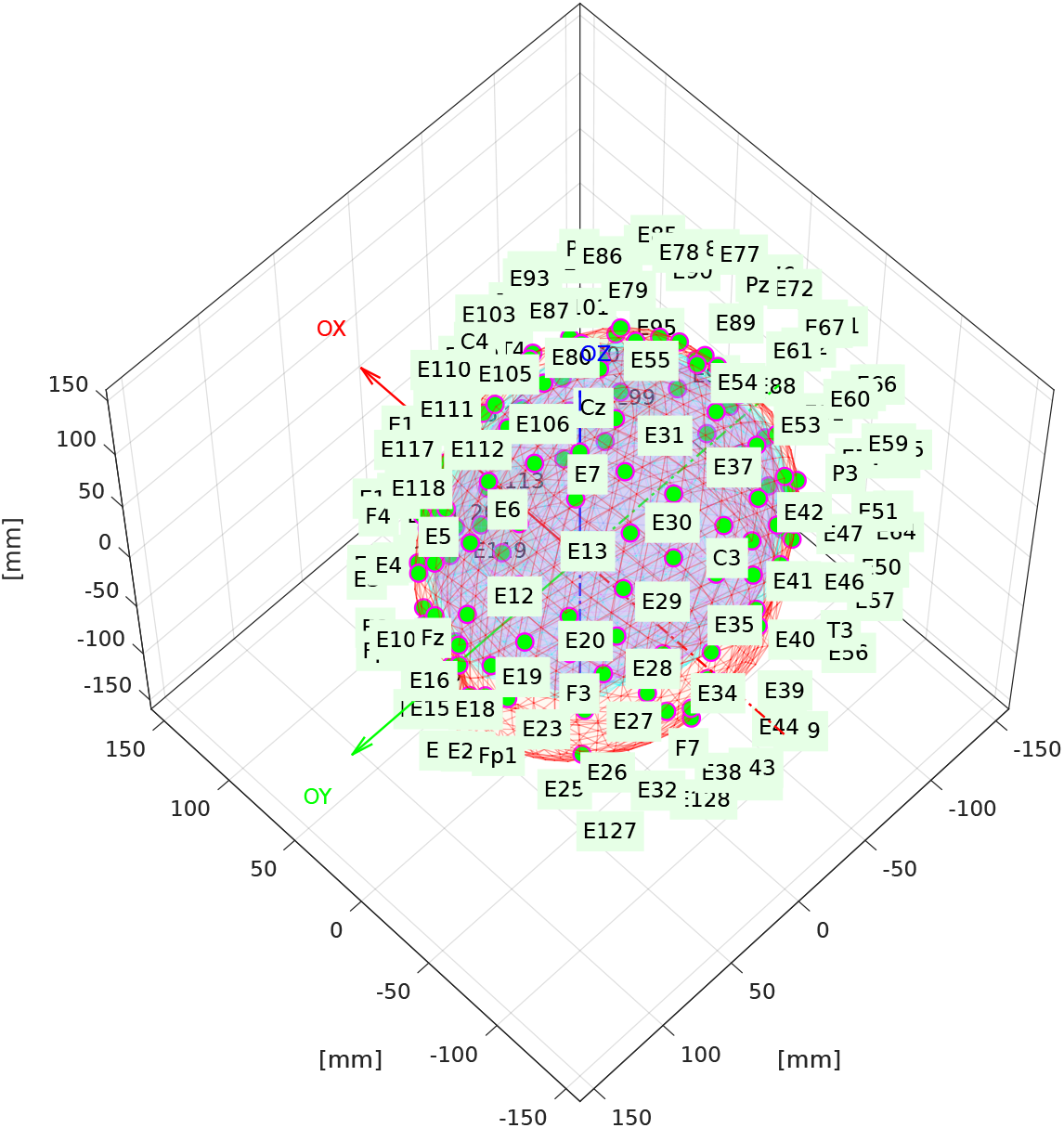
Volume conduction model essential components. Triangulation meshes representing brain, skull and scalp boundaries with electrode positions plotted on top of the scalp surface. This figure was generated using an instance of `EEGPlotting` class employing `plotbrainoutermesh(), plotskulloutermesh(), plotscalpoutermesh(), plotelectrodepositioning()` and `plotelectrodelabels()` methods.

**Fig. 7.**
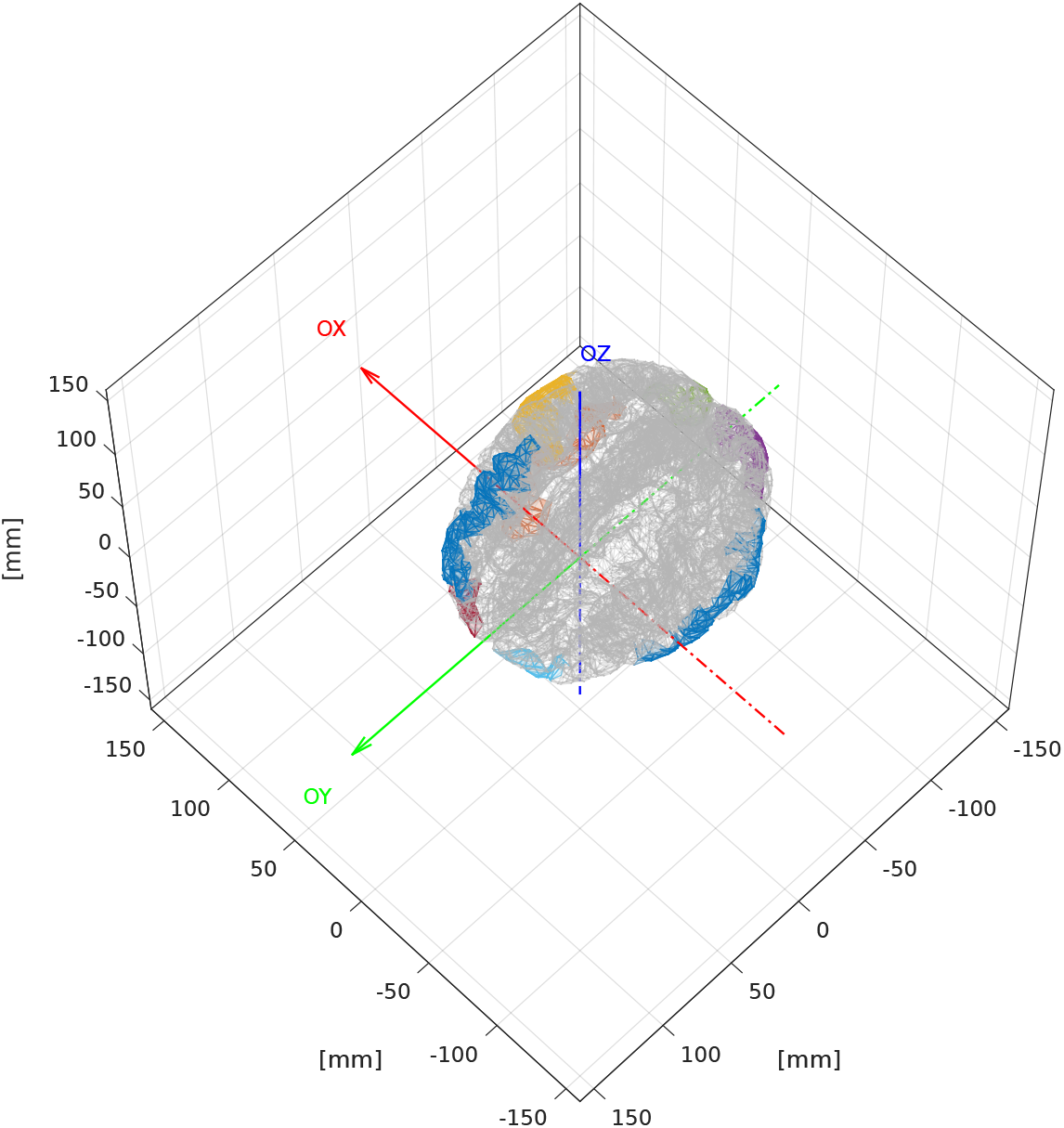
Cortex and ROIs. Detailed cortical surface triangulation with selected cortical patches. This figure was generated using an instance of `EEGPlotting` class employing textttplotcortexmesh() and textttplotROIvisualization() methods.

**Fig. 8.**
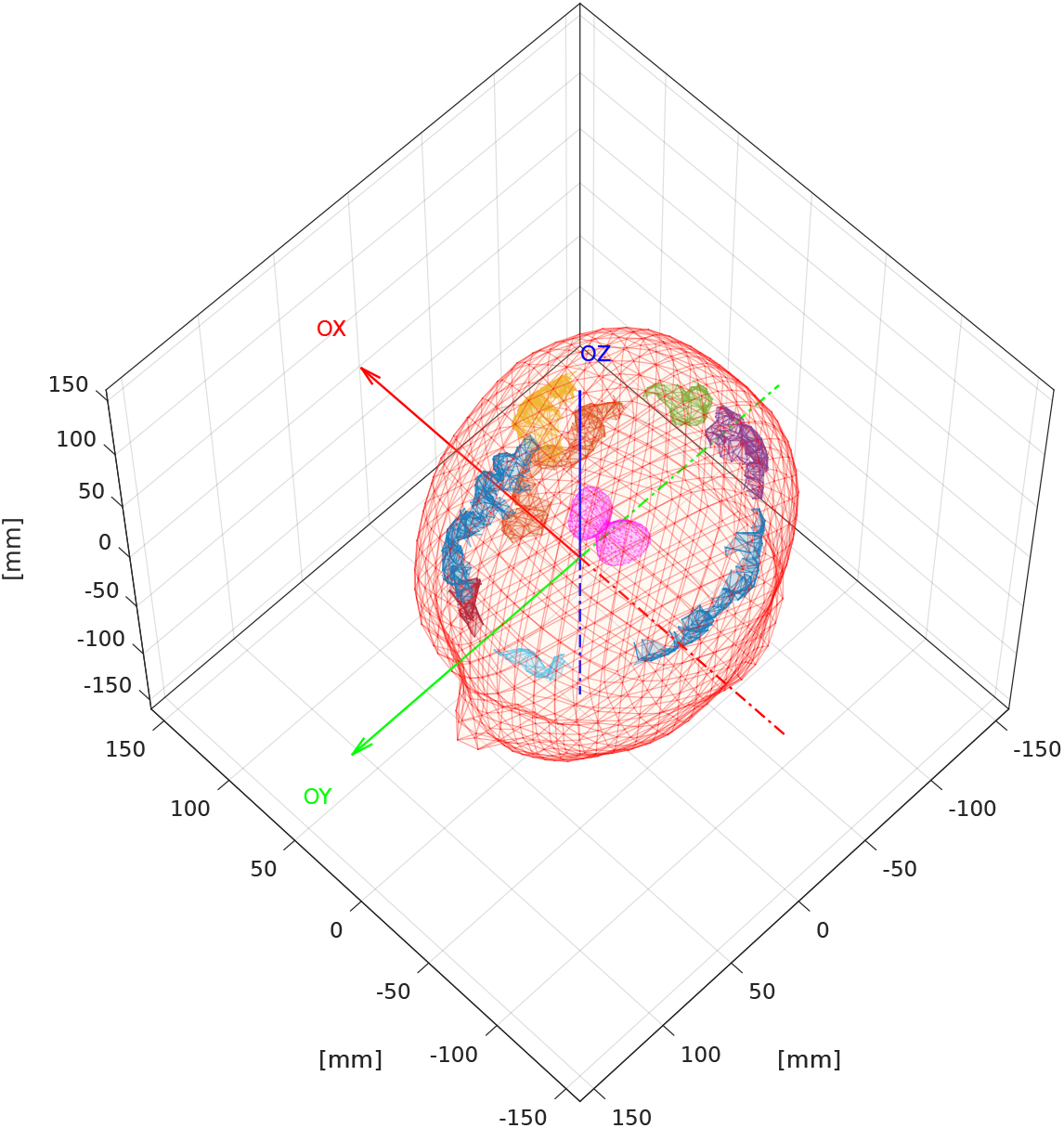
ROIs and thalami. Cortical patches selected as a candidate ROIs for source position with thalami mesh and scalp outer mesh. This figure was generated using an instance of `EEGPlotting` class employing `plotROIvisualization(), plotdeepsourcesasthalami()` and `plotscalpoutermesh()` methods.

**Fig. 9.**
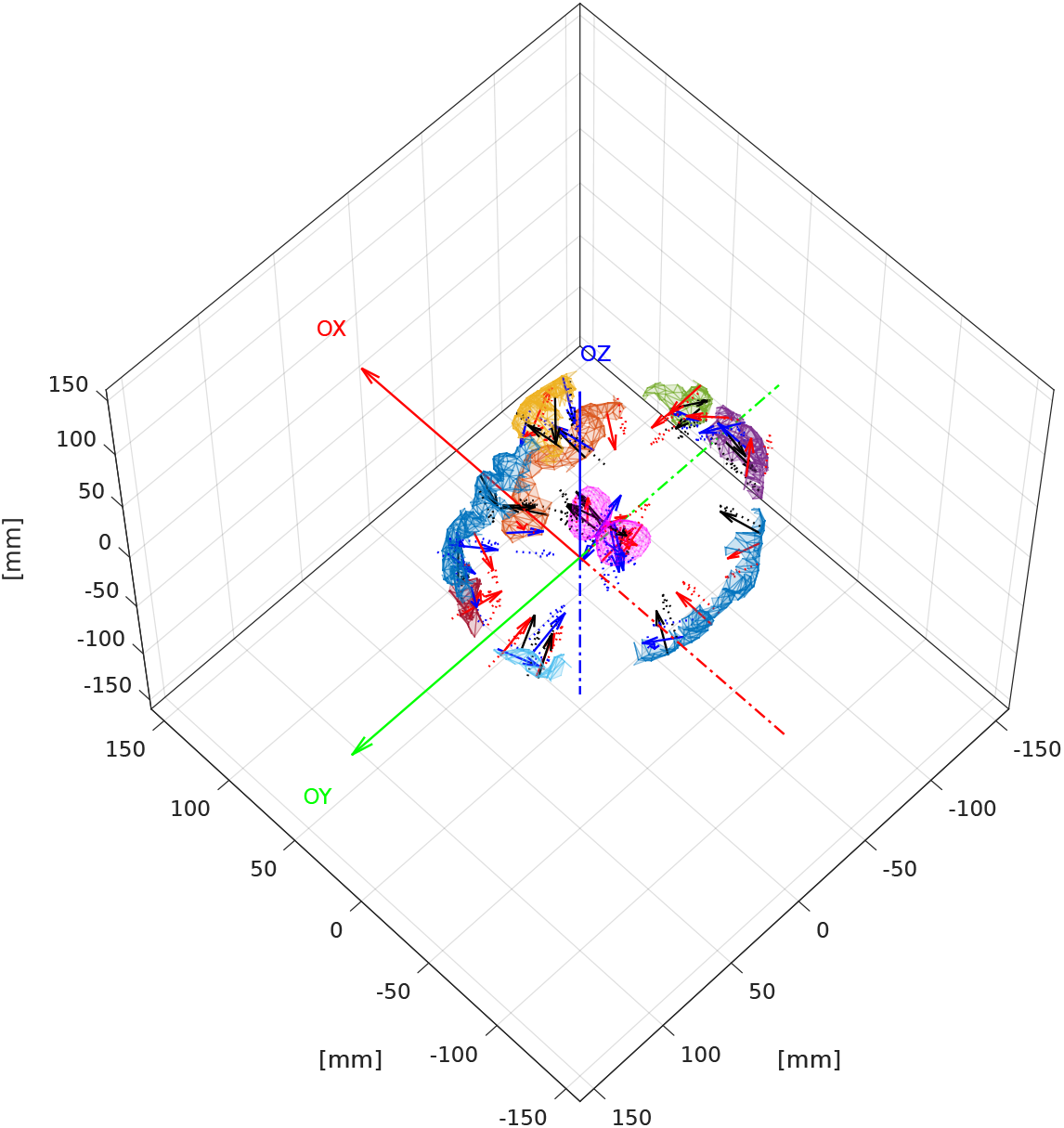
Bioelectrical activity positions and orientations. Cortical patches selected as a candidate ROIs for source position with thalami mesh. Vectors representing direction of the dipole moments for the sources of bio-electrical activity. Red lines represent the activity of interest; blue — the interfering activity and black — background activity. Solid lines represent original sources and dotted lines represent perturbed sources. Arrows representing dipole position and orientation are drawn not to scale. This figure was generated using an instance of `EEGPlotting` class employing `plotROIvisualization(), plotdeepsourcesasthalami()` and `plotsourcevisualization()` methods.

Plot consists of layers that are generated by functions with self-explonatory names. E.g. function `plotROIvisualization` plots cortex, regions of interest. Function plotsourcevisualization plots mesh for ROIs on cortex, mesh for deep sources ROI, sources and cortex.

– `EEGPlotting:`

– `plotMVARmodelcoefficientmatrixmask` — this method plots mask for MVAR model coefficient matrix.
– `plotPDCgraph` — this method is used for plotting PDC profiles across sources of interest and interfering sources.
– `plotDTFgraph` — this method is used for plotting DTF profiles across sources of interest and interfering sources.
– `plotMVARmodelcoefficientmatrixgraph` — this method plots the matrix of composite MVAR model for sources of interest, interfering and background sources.
– `ploterrortable` — this method plots results of reconstruction as heatmap table.
– `plotdeepsourcesasicosahedron642` — this method is for plotting of deep sources.
– `plotdeepsourcesasthalami` — this method can be used for plotting of both thalami.
– `plotcortexmesh` — this method plots cortex mesh.
– `plotbrainoutermesh` — this method plots brain outer mesh.
– `plotskulloutermesh` — this method plots skull outer mesh.
– `plotscalpoutermesh` — this method plots scalp outer mesh.
– `plotelectrodepositioning` — this method plots electrode positions.
– `plotelectrodelabels` — this method plots electrode labels.
– `plotROivisualization` — this method plots ROIs based on generated meshes.
– `plotsourcevisualization` — this method is used for source visualization.

### 6.4 Unit test class

Since this implementation is based on the previous one, which was done in `Org-mode`, authors have created a class `EEGTest` for unit tests.

In order to generate and distribute files into directories (necessary for the test) use a `make` target `test`.

For example, unit tests can look like that:

~~~
1 eegtest = EEGTest()
2 eegtest = eegtest.testsetup()
3 eegtest = eegtest.testsignals()
4 eegtest = eegtest.testleadfields()
5 eegtest = eegtest.testfilters()
6 eegtest = eegtest.testerrors()
~~~

This class can be useful for developers who wants to extend our framework. This way they always can check whether it still passes the compliance test.

## 7 Sample usage

### 7.1 Generating set of parameters for simulations

Generation of a new simulation requires preparation of a set of parameters. This is done by the `EEGParameters` class, in which the `generate` method is included. In the default version the `dummy` function is called, which returns the default parameter structure, but the user can always overwrite it to create her/his own version, or change the configuration of the simulation and perform own experiments. Syntax for generating parameters is as follows:

~~~
1 parameters = EEGParameters().generate();
~~~

Several sample settings have been prepared. Functions `setinitialvalues` and `setsnrvalues` will set up initial parameters and signal to noise ratios. Function `smartparameters` overwrites initial values and contains parameters for the sample run. Function `testparameters` contains parameters for unit test done by `EEGTest`. To use it line containing this function must be uncommented.

Generated structure contains about 50 fields. All fields are listed in Table 1.

### 7.2 Example run of simulations

The input configuration structure from `EEGParameters` class contains options and parameters that specify how the stimulation will run. Once the user is satisfied with the parameter settings a sequence of simulations is ready to run, which may look, e.g., like that:

~~~
 1 filters = [‘LCMV’, ‘MMSE’, ‘ZF’, ‘RANDN’, ‘sMVP_R’];
 2 reconstruction = EEGReconstruction();
 3 reconstruction = reconstruction.init();
 4 for np = 1:length(parameters)
 5  parameter = parameters(np)
 6  reconstruction = reconstruction.setparameters(parameter);
 7  reconstruction = reconstruction.setsignals();
 8  reconstruction = reconstruction.setleadfields();
 9  reconstruction = reconstruction.setpreparations();
10  reconstruction = reconstruction.setfilters(filters);
11  reconstruction.save();
12 end
13 reconstruction = reconstruction.printaverageresults();
~~~

Here `parameters` were generated as described above. Selection of filters to compute the source reconstruction is done above in a very direct way. Filters are self-contained, i.e. they can run independently. What is worth noting, there are fifteen spatial filters available in the current version of supFunSim (including, e.g., classical LCMV), and more will be added.

All intermediate values of model variables along with the initial settings are stored in attributes `SETUP` and `MODEL`. Attribute `RESULTS` contains the scores of all the most important measures of errors for individual filters. Moreover, all meshes are kept in attribute `MATS`. The main reason is that meshes are loaded only once during all iterations. All that, as a part of the main object, can be saved in mat file and later restored.

~~~
1 load(‘reconstruction_DATE.mat’);
2 obj.MODEL
~~~

Here `DATE` is identifier of reconstruction, given by date of execution.

## 8 Conclusion

Many tools have been created for EEG/MEG source reconstruction and localization. To the best of our knowledge our supFunSim toolbox is the first written in modular and object-oriented way that allows to use many spatial filters for EEG source reconstruction. Except for classical linearly constrained minimum-variance (LCMV) filter, [10,38,36], eigenspace LCMV [36], nulling (NL) [16], it also contains our new minimum-variance pseudo-unbiased reduced-rank (MV-PURE) filters [33,30,32]. Our toolbox may thus enable an easy comparison of accuracy of different filters. It may be used in combination with the standard FieldTrip or EEGLab packages, and should be useful in construction of both forward and inverse models. The use of reconstructed signals for the directed connectivity analysis using partial directed coherence (PDC) [2] and directed transfer function (DTF) [18] measures is also of great interest in analysis of information flow in the brain.

Future work will proceed on the further development of supFunSim toolbox, and on the testing of the methods on real data from EEG experiments. Many large-scale subnetworks have been discovered using fMRI, and a big challenge is to characterize their dynamics. EEG signals after source reconstruction may help to analyze fast switching between different subnetworks, discovering fingerprints of brain’s cognitive activity.

Matlab is still a very popular language among neuroscientists and it was an obvious choice due to availability of functions that could be adopted from other toolboxes. However, we plan to make a Python implementation and thorough simulations illustrating the power of our toolbox, comparing different filters. The code in the current form that contains objects should be easily rewritten to Python.

## A Appendices

### A.1 Implemented spatial filters

We denote the concatenated composite lead-field matrices of *H* and 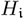 as 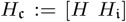, and similarly 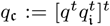. The covariance matrices of *q*, 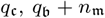, *y* are denoted by *Q*, 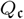, *N, Y*, respectively.

We selected for comparison the following spatial filters:

1. The LCMV filter, expressed as both [23]

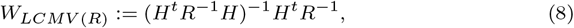

and [22]

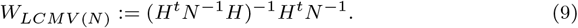
2. The nulling filter [16]:

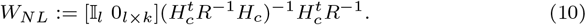
3. The Wiener filter, defined as [17]

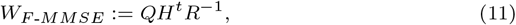

for the interference-free model, and [17]

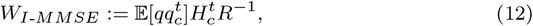

for the model in presence of interference.
4. The zero-forcing filter, defined as [17]

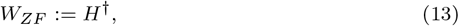

where *H*^†^ denotes pseudo-inverse of *H*.
5. The eigenspace-LCMV filters [36] exploiting projection of the signal covariance matrix *R* onto its principal subspace of the forms

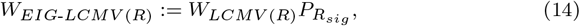

and

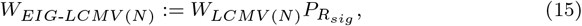

where *P_R_sig__* is the orthogonal projection matrix onto subspace spanned by eigenvectors corresponding to λ_1_ ≥ ⋯ λ_*sig*_ — the *sig* largest eigenvalues of *R*, where *sig* is the dimension of signal subspace.
6. The MV-PURE filters, defined as [32]

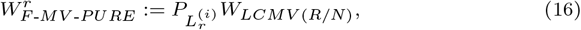

for the interference-free model, and [32]

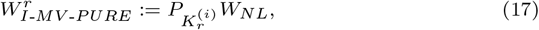

for the model in presence of interference. In the above expressions, 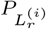, for *i* = 1, 2, 3, are the orthogonal projection matrices onto subspaces spanned by eigenvectors corresponding to the *r* smallest eigenvalues of symmetric matrices

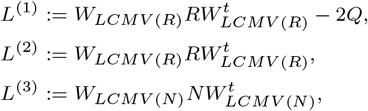

respectively; similarly, 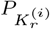, for *i* = 1, 2, 3, are the orthogonal projection matrices onto subspaces spanned by eigenvectors corresponding to the *r* smallest eigenvalues of symmetric matrices

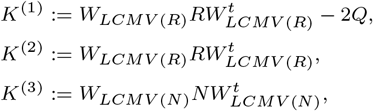

respectively. Here, *Q* is the covariance matrix of sources of interest *q*. The file `EEGReconstruction.ipynb` is richly commented using mathematical formulas, thanks to which it will be easy to find a concrete place of implementation of a specific filter.

### A.2 Configuration

Table 1 contains all configuration parameters. Some of the more complex parameters have been explained below the table.

Within `SETUP`, perhaps the most important are the parameters controlling configuration of activity of sources (`SRCS`), configuration of deep sources (`DEEP`), number of samples and realizations of the signal (`n00, K00`), presence of ERPs (`ERPs`), number of iterations (`ITER`), configuration of lead-fields perturbation (`CUBE, CONE`), signal to noise ratios, (`SINR, SBNR, SMNR`), and presence of signal components (`SigPre, IntPre, BcgPre, MesPre, SigPst, IntPst, BcgPst, MesPst`).

There are 7 structures containing head conduction model. They are listed in Table 2.

**Table 2:**
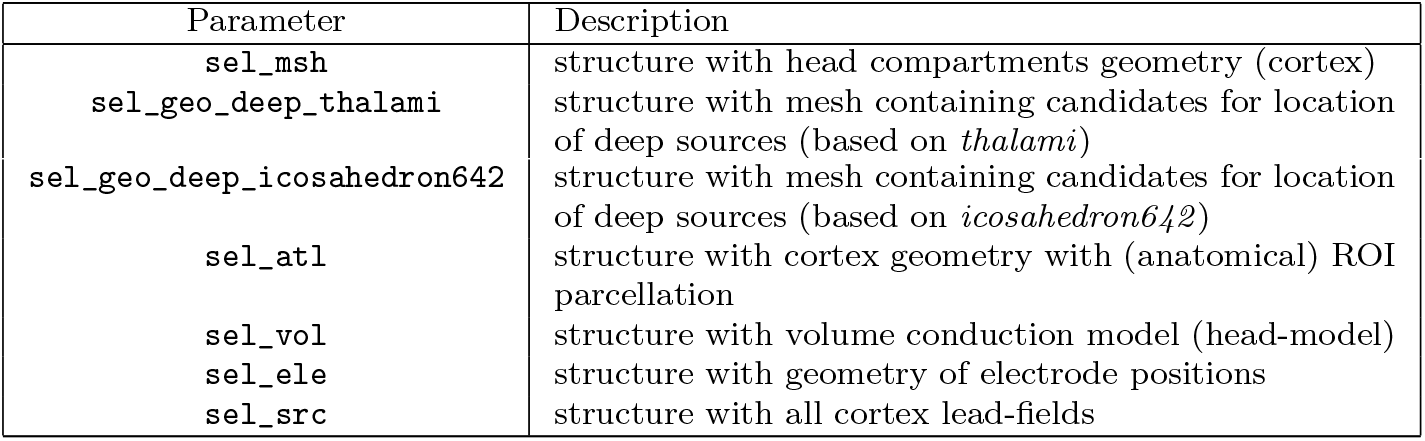
`MATS` structure containing all meshes.

| Parameter | Description |
| --- | --- |
| **sel_msh** | structure with head compartments geometry (cortex) |
| **sel_geo_deep_thalami** | structure with mesh containing candidates for location of deep sources (based on *thalami*) |
| `sel_geo_deep_icosahedron642` | structure with mesh containing candidates for location of deep sources (based on *icosahedron642*) |
| `sel_atl` | structure with cortex geometry with (anatomical) ROI parcellation |
| `sel_vol` | structure with volume conduction model (head-model) |
| `sel_ele` | structure with geometry of electrode positions |
| `sel_src` | structure with all cortex lead-fields |

## Acknowledgements

This study was supported by the National Science Center Poland, grant no. UMO-2016/20/W/NZ4/00354.

## Footnotes

1 Spatial filters are also known as beamformers in array signal processing.

2 > “*The great topmost sheet of the mass, that where hardly a light had twinkled or moved, becomes now a sparkling field of rhythmic flashing points with trains of traveling sparks hurrying hither and thither. The brain is waking and with it the mind is returning. It is as if the Milky Way entered upon some cosmic dance. Swiftly the head mass becomes an enchanted loom where millions of flashing shuttles weave a dissolving pattern, always a meaningful pattern though never an abiding one; a shifting harmony of subpatterns*”. > > — – Charles S. Sherrington, Man on his nature. 1942

3 It can be download from http://neuroimage.usc.edu/bst/download.php?file=tempColin27_2012.zip and contains: `tess_innerskull.mat`, `tess_outerskull.mat` and `tess_head.mat`.

4 Available at http://www.fieldtriptoolbox.org/tutorial/headmodel_eeg_fem/

